# Trait-environment relationships are predictive, but not general across species

**DOI:** 10.1101/2020.01.13.904698

**Authors:** Dachin N. Frances, Amelia J. Barber, Caroline M. Tucker

## Abstract

Understanding the relationships between organisms and their environments is increasingly important given human impacts on global conditions. However, predicting how community diversity and composition will change in the future remains challenging (Mouquet *et al* 2015). One recent approach is to use traits to mechanistically inform how environmental conditions affect performance (i.e., trait-environment relationships), under the assumptions that these measures relate to each other in predictive and general ways. Unfortunately, results have been inconsistent, ignore phenotypic plasticity, and rely heavily on observational data (Shipley *et al* 2016). We evaluated the predictability and generality of trait-environment relationships in a controlled experimental microcosm system of four daphniid species. We cultured each species along a stressful gradient (conspecific density), measuring performance (fecundity) and traits related to performance (body length, 2nd antenna length, eye diameter, relative growth rate, and age at first reproduction). Using structural equation models, we evaluated the role of traits in mediating changes in individual fecundity in response to conspecific density. We built models for each species separately considering within-species trait variation, and for all species together by considering all trait variation across the four species. Results from this controlled system highlight that the relationship between individual traits and the environment (conspecific density) is strong and predictive of performance (fecundity), both within- and across-species. However, the specific trait-environment relationships which predicted fecundity differed for each species and differed from the relationships observed in the interspecific model, suggesting a lack of generality. These results will inform and improve the use of traits as a tool for predicting how changing environments will impact species abundances and distributions.

## Introduction

The responses of individuals, populations, and communities to various environmental gradients is of long-standing interest in ecology [1] and evolution [2]. In an unprecedented era of rapid anthropogenic change, the ability to describe and predict these responses to rapidly changing biotic and abiotic conditions is imperative for conservation, restoration, and management [3]. Recent work [4,5] has highlighted the potential to use traits — measurable characteristics that describe the phenotype, such as morphology, physiology, phenology, and behaviour [6] — to link environmental conditions and individual performance. Logically, if individual phenotypes reflect adaptive selection for success in a particular environment [7], then traits should predictably relate to performance in different environments (i.e., trait-environment relationship) [8,9]. Some studies have reported moderate to strong correlations between traits and environmental conditions, particularly at large spatial scales and for broad climatic gradients like temperature and moisture [10]. On the other hand, studies frequently fail to identify significant or predictive relationships between traits and the environment [9,11,12]. Drawing general conclusions is complicated by the large range of spatial scales, species, and methodological approaches considered by researchers. This complexity must be addressed, to determine if and how trait-environment relationships can be used to make predictions about the responses of ecological systems to environmental change. Addressing an independent but related set of questions should clarify our understanding of trait-environment relationships: What are the functional forms of trait-environment interactions (i.e., if performance is predicted with a function relating traits and the environment, what is the function)? Given the functional form, how much variation in performance does the relationship explain (how predictive is the relationship)? Are these functional relationships comparable, either between species, and/or across taxonomic groups (how general is the relationship)?

Identifying predictive and general trait-environment relationships has not proven to be straightforward. One problem is that it is difficult to tease apart evidence that these relationships are not predictive or general across species from the inherent difficulties in measuring trait-environment relationships. Studies (frequently on plants) of traits and environmental gradients most often rely on observational data collected at a variety of spatial scales. The presence of an individual at any given point along a gradient is influenced by multiple processes beyond the abiotic environment, including dispersal limitation, competition, predation, and mutualisms, all of which can distort estimates of the underlying trait-environment relationships [13]. Methodological issues with inferring these relationships from observational data are well-known. The Fourth Corner problem, for example, refers to the inherent difficulty in quantifying the strength of trait-environment associations if they are inferred indirectly (see [14]). Analyses of observational data can also differ greatly in terms of the type of trait data available – some may quantify trait values directly (field measurements) but it is also common to obtain measures through the use of large databases (e.g. TRY [15]). Despite the often implicit assumption that trait values should reflect selection for specific phenotypes in particular environments, they frequently also reflect plastic responses to the environment [16] and this plasticity may play an important, yet under-considered role in determining performance. Finally, measures of performance are often approximations of fitness such as growth, reproduction, survival, or dispersal [8,9,12] which may be imperfect proxies [6,9], or only provide short-term estimates of performance, especially in long-lived species.

In addition to the requirement that trait-environment relationships be predictive, they should also be general across multiple species and/or taxonomic scales. If an optimal trait value does exist for a particular environment, then the taxonomic scale at which the trait is measured (among-individuals, among-populations, among-species, among-communities) should be irrelevant [8,9]. For instance, Vasseur *et al* (2012) showed that there was sufficient variation and similar tradeoffs within *Arabidopsis thaliana* to produce an intraspecific leaf economics spectrum [17]. This type of generality has been identified in at least some studies of individuals, populations, and species [18,19]. But other studies hint at potential inconsistencies in trait-environment relationships both when compared between species [20] or across ecological or spatial scales [21,22].

Addressing fundamental questions about trait-environment relationships – about the functional form of these relationships, the predictive ability of this functional relationship, and the generality of the relationship – is well-suited to work in highly controlled, replicable experimental systems with short generation times. We use a novel microcosm system containing freshwater daphniid species (*Daphnia magna, Moina micrura, Simocephalus vetulus, Ceriodaphnia dubia*). These species are ecologically important consumers and prey, have rapid generation times and simple morphologies, and many candidate functional traits [23,24]. Based on the wealth of information on their ecology and life histories [25,26], development and genetics (e.g., https://genome.jgi.doe.gov), and theoretical ecological models [27], they are excellent model organisms for ecology and evolution [28]. Using this system, we ask two key questions. 1) Are there functional relationships between an individual’s traits and their performance along an environmental gradient, and how much variation is explained by this relationship? 2) Are these functional relationships the same for different daphniid species, or when modelled at different taxonomic scales (e.g. within- and between-species)? Describing the functional form of these trait-environment relationships is also in need of further study, but optimally, functional relationships should be mechanistic and built from fundamental biological principles rather than statistical or correlational relationships (e.g. see [29,30]). Such an approach is outside of the scope of this study, but work with daphniids has begun to address this question [27]. For the purposes of this study, we estimate these functional relationships statistically using first order approximations.

To address these questions, we experimentally manipulated environmental conditions, measured individual trait values and quantified fecundity. For each species, we varied the density of conspecific individuals in a given microcosm from 1, 2, 4, 6, to 8 individuals, creating an increasingly stressful biotic gradient. We expected this gradient to have negative effects on individual growth, survival, and reproduction. High conspecific densities are associated with reductions in per capita resource availability as well as crowding, which can lead to changes in feeding behaviour and reproduction [31,32]. Each day we observed the focal generation’s development, reproduction (counting and removing all juveniles born), and measured fecundity (the total juveniles produced per adult per microcosm). At senescence, we measured ecologically relevant traits related to reproduction (body length and age at first reproduction [33]), feeding (length of 2nd antenna [25,34]), and energy allocation (eye diameter and relative growth rate [23,35,36]; see Supplementary Materials Table S1 for references and justification of traits). To explore whether there were functional relationships between the traits and the conspecific density treatment on performance, and further, if they are predictive or general, we used structural equation modelling (SEM). This modelling approach allowed us to tease apart the direct and indirect (through trait changes) impacts of conspecific density on fecundity and allowed the hierarchical structure of the data to be explicitly modeled [37]. We fit the same model structure in all cases – one that included all potential direct paths between density, trait values, and fecundity. We fit this model for two taxonomic scales (describing within-species or among-species trait-environment relationships). This allowed us to test whether there are significant trait-environment relationships, and if so, to ask how well they predict fecundity. We also tested whether these relationships are general in form for all species, or when compared across the two taxonomic scales. We considered a trait-environment relationship as significant when we could identify traits which were both significantly impacted by density, which were also significant predictors of fecundity.

## Results

### Effect of density on raw trait values

Increasing conspecific densities led to notable changes in individual phenotypes, including morphological traits (body length, 2nd antenna length, and eye diameter; Figure 1a-c). The extent of these changes appeared to be species-specific, although the direction of change was generally consistent, with higher densities leading to smaller average body lengths, eye diameters, and 2nd antenna lengths. Similarly, relative growth rate (RGR) decreased with density (except *C. dubia*; Fig. 1d). Variation in trait values tended to decrease with density (e.g., *D. magna, S. vetulus, and C. dubia* for eye diameter; Fig. 1).

**Figure 1.**
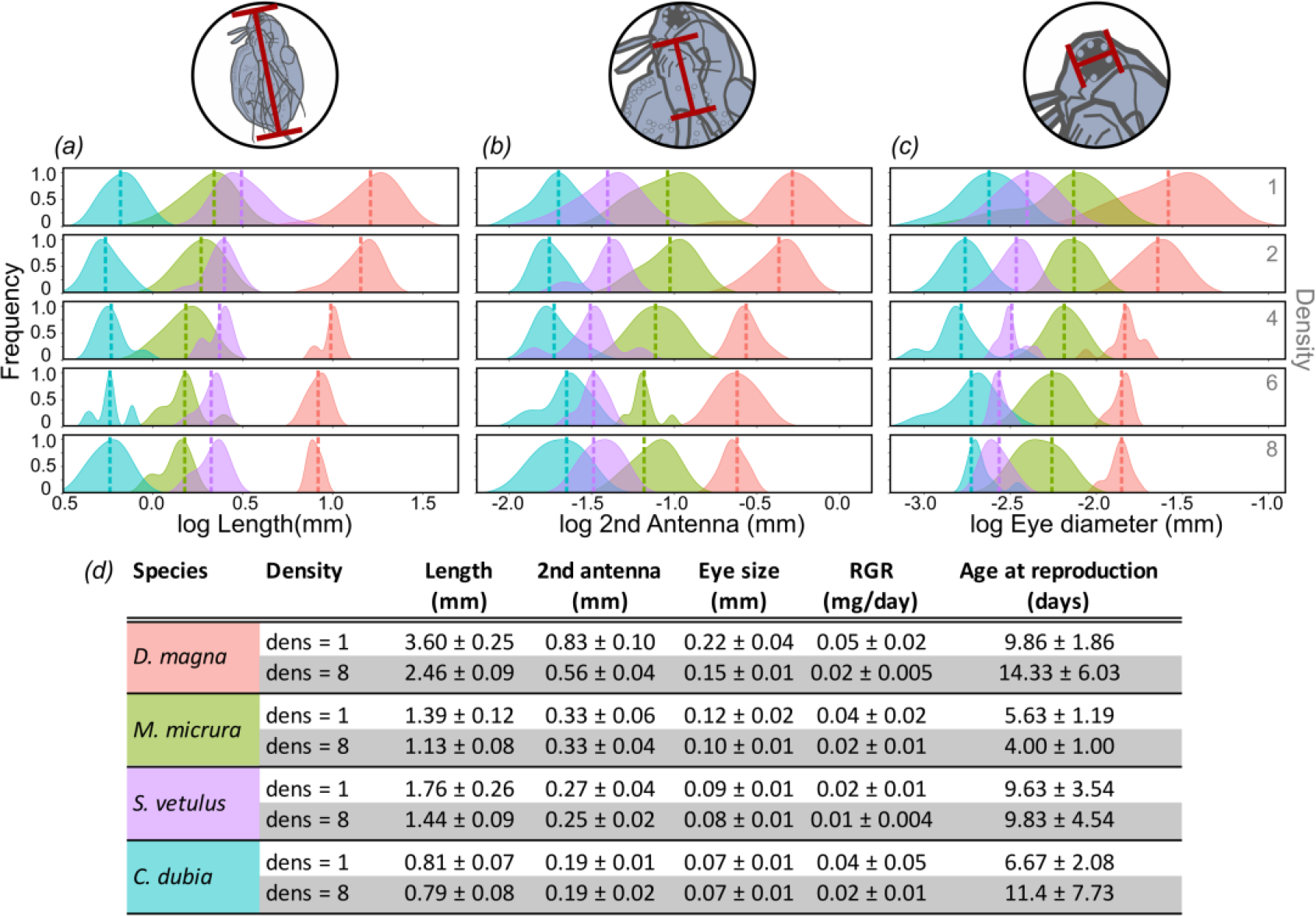
Summary results for the effects of conspecific density on species’ observed trait measures. *a*-*c)* Changes in the frequency distributions of the three morphological traits (body length, 2nd antenna, eye diameter) with increases in density (from top row, density = 1, to the bottom row, density = 8). Dashed vertical lines are the average trait value per density treatment. *d)* Summary of changes in all traits, per species, specifically comparing the trait value at the lowest density to that at the highest density. Values are means ± SD.

### Intraspecific models of trait-environment interactions for daphniid species

#### Model prediction & generality between species

We fit a structural equation model to the individual data collected per species. The intraspecific SEMs explained the majority of the variation in individual fecundity for a species, as a combination of indirect effects of density on fecundity via traits (all species except *C. dubia*) and direct relationships between trait values and fecundity (all species, see Fig. 2). For *D. magna*, relative growth rate and age at first reproduction were significantly affected by density (52% of the variation in RGR is explained by density and 11% of the variation in age at first reproduction), and significant predictors of fecundity (*p* < 0.001; whole-model R2 for fecundity was 0.95; Fig. 2). For *S. vetulus and M. micrura* significant relationships between density and fecundity were mediated by body length (R2 = 0.20 and 0.22 respectively), eye diameter (R2 = 0.18 and 0.12 respectively), and relative growth rate (R2 = 0.11 and 0.09 respectively); model R2 for fecundity were 0.83 and 0.62, respectively. Notably, we did not find significant trait-environmental relationships for *C. dubia*, although density had a marginal effect on the age at first reproduction (for the path between density and age at first reproduction, *p* = 0.0589).

**Figure 2.**
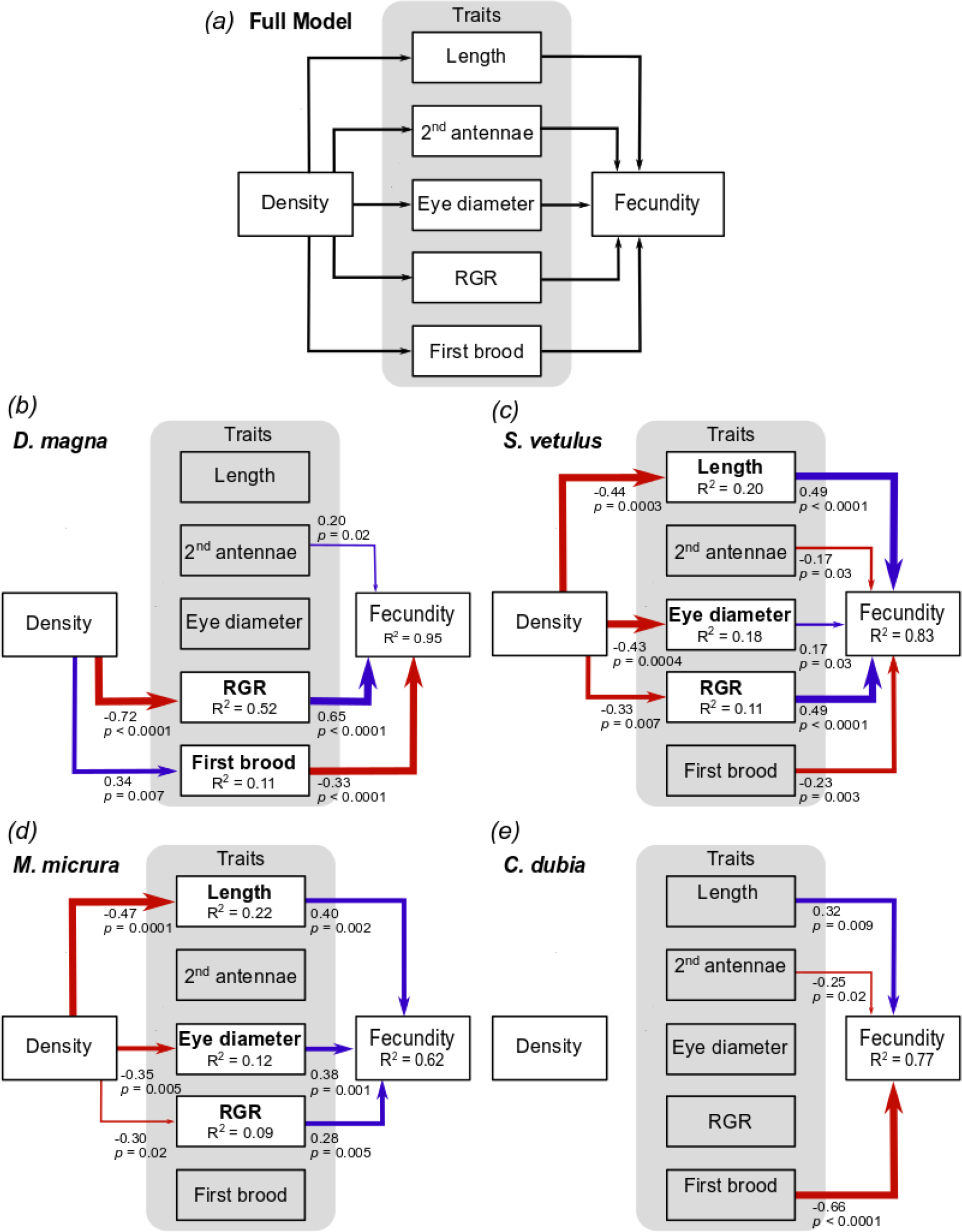
Results for the intraspecific structural equation models. *a)* Shows the full model structure fit separately to each species’ data. This assumes that all potential paths between conspecific density and fecundity are possible. *b*-*e)* show the set of significant relationships for *b) D. magna, c) S. vetulus, d) M. micrura*, and *e) C. dubia*. Arrows represent the standardized path coefficients; associated information includes coefficient value and significance. These are also shown visually: arrow width is scaled with significance of the coefficient (wider arrows have smaller *p*-values), and blue arrows identify positive path coefficients while red arrows signify negative path coefficients. R2 values for each linear model are also shown

When the structure of these intraspecific models was compared across species, there was a lack of generality in terms of which traits were significant for each species (although some of the same traits were significant for several species). Though significant paths varied between species, it is worth noting that the direction of observed trait-environment relationships was consistent across species. Increasing densities always led to smaller morphological traits, slower relative growth rates, and older ages for the onset of reproduction (Fig. 2 b-e). Model fit to observed data is shown in the Supplementary Materials Fig S1.

### Interspecific models of trait-environment relationships for daphniid species

#### Model prediction & generality across taxonomic scales

We also asked whether an interspecific SEM which combined traits from all species could identify general trait-environment relationships that describe fecundity across all four species. The SEM containing all species data identified three key traits which were significantly impacted by density (relative growth rate, R2 =0.11, body length, R2 =0.10, age at first reproduction, R2 =0.03), and also have significant impacts on fecundity (R2 for fecundity was 0.72; Fig. 3). The good fit of the model’s predicted values to the observed data confirms is shown in Fig. 3. This interspecific scale model appears to identify general (multi-species) patterns in trait-environment relationships. We also confirmed, using a multivariate ANOVA, that the multivariate trait values followed a similar trajectory in response to density, regardless of species. The shift in the mean trait values was significantly predicted by both the density treatment and species identity (*p* < 0.01, Supplementary Material, Table S3), but the interaction term was small and non-significant (marginal at *p* = 0.05119).

**Figure 3.**
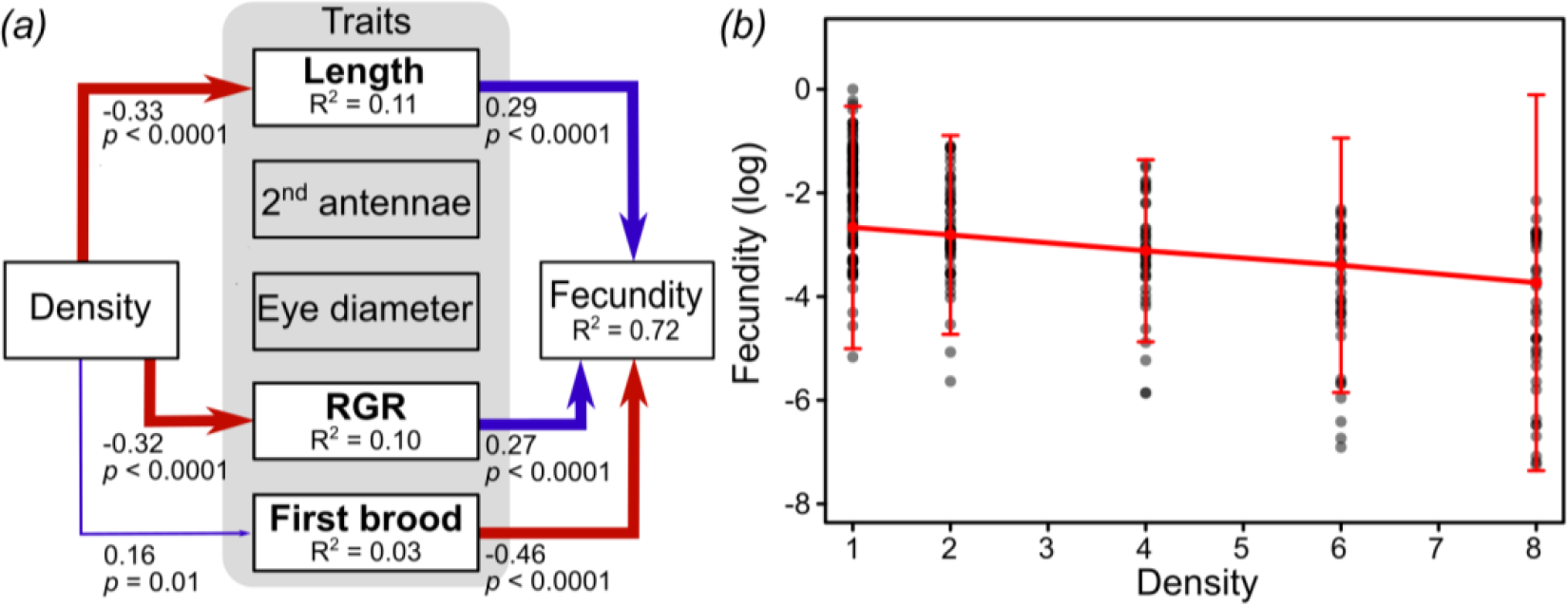
*(a)* Results for the interspecific structural equation model, calculated using trait values from all four species combined (*D. magna, S. vetulus, M. micrura*, and *C. dubia*). Arrows represent the standardized path coefficients; associated information includes standardized coefficient values and significance. Arrow width represents the significance of the path coefficient (wider arrows have smaller *p*-values), and blue arrows identify positive path coefficients while red arrows signify negative path coefficients. R2 values for each linear model are also shown. *(b)* Model fit, showing predicted fecundity, in red, per density treatment. We calculated the mean and standard error of each trait for a given density treatment level and used these to define a normal distribution for each trait combination. We drew randomly from these distributions and used the trait values to calculate fecundity. This procedure was repeated 1000 times for each density. Grey points are the observed values of fecundity. Fecundity is a rate of juveniles produced per adult per day, scaled by the max fecundity across species, and then log-transformed. Error bars are 95% confidence intervals

## Discussion

One goal of research into functional relationships between environmental conditions and organism performance is the development of predictive trait-based models of species’ distributions and community composition (e.g. [38,39]) Recent works have highlighted limitations in achieving this goal, e.g., traits often appear to have only weak or absent relationships with performance [9]. In this study, we establish tests of the underlying assumptions for trait-environment relationships, including that they should be predictable (that is, variation in trait values in relation to the environment explain significant variation in performance) and general (that is, functional relationships are of similar form among species or across taxonomic scales) in a controlled, experimental system. We found that traits are predictive of fecundity and the structure of these trait-environment relationships is general across the four species, but also that these results are context-dependent in interesting and informative ways. Specifically, we found that a combination of five functional traits (relative growth rate, age of first reproduction, body length, 2nd antenna length, and eye diameter) explained the majority of variation in fecundity across a stressful biotic gradient. However, these models of intraspecific trait variation were not always predictive; for one species (*C. dubia*) no traits were significant predictors. Further, the other intraspecific models were not general and differed among species in terms of which traits were significant predictors of fecundity. In contrast, our model of interspecific trait differences was both general and predictive, suggesting traits alone can be sufficient to describe responses to the density gradient.

For all intraspecific models, we found that the five traits measured here explained significant variation in individual fecundity, highlighting the utility of traits for describing performance. The traits we selected have known relationships with daphniid ecology and life-history (e.g., [23,24], see Table S1 for details). For example, body length is associated with growth rate, filtering rate, and survival, and thus with fitness [40]. RGR is directly related to resource and energy allocation towards growth [23]. Age at first reproduction is a trait that commonly trades off with lifespan [33] and in daphniids, earlier maturation is correlated with higher intrinsic rates of increase [41]. For all species except *C. dubia*, we identified significant axes of trait variation in response to the environment, specifically related to the pace of life history and body size which are common responses across multiple groups of species [42]. With increasing stress, fecundity decreased as a result of declines in body size (length, eye diameter), and/or shifts towards slower growth rates and delayed onset of reproduction. As shown in Supplementary Fig. S1, model predictions of fecundity were accurate (and the corresponding R2 values were high), likely aided by the high temporal resolution of sampling, and the ability of the microcosm environment to be carefully controlled, including the age and maternal background of starting individuals as well as non-focal abiotic conditions.

Notably, we found no significant trait-environment relationships for *C. dubia*, although there was a trend of slower age to first reproduction at higher densities (*p* = 0.0589), and this trait was significantly (negatively) associated with total individual fecundity (*p* < 0.0001). There are a number of reasons we may not have identified significant predictors of *C. dubia*’s performance along the conspecific density gradient. Because of the size differences among the four species, one explanation is perhaps the smaller species (*M. micrura* and *C. dubia*), experienced the crowding aspect of the density gradient more weakly than larger species (*D. magna* and *S. vetulus*). We performed an additional experiment with a small (*M. micrura*) and large (*D. magna*) species, where we manipulated food availability rather than conspecific density. This let us match the gradient of food availability with that in the original experiment, but we fixed the number of individuals to one. Crowding did not impact the large species (*D. magna*) more than the small, in fact we found that there was no meaningful difference in the results (i.e., fecundity) from these experiments, for either of the two species (*p* = 0.49 and *p* = 0.83 for *M. micrura* and *D. magna*, respectively; Supplementary Materials Table S4, Fig. S2). This suggests that food limitation is the primary driver of our results.

When modelling within-species trait variation, we found that even among these four ecologically similar species, there was no general combination of traits that predicted fecundity. Trait correlations can cause non-independent responses of traits, and if these correlations are different within-species, they may constrain individual responses and make generality less likely. Such differences in trait correlations may reflect underlying developmental, physiological, evolutionary, ecological, and genetic constraints, and therefore selective pressures [43]. Among-species, and at higher taxonomic scales, consistency in trait correlations is perhaps more likely [44], perhaps explaining the greater generality for the among-species model. The trait differences among species, related to the pace of development (RGR, age at first reproduction) and body size (length) responded to the density gradient in a predictable and general fashion. Not only did interspecific trait values mediate interactions between the environment and density, they explained 72% of the overall variation in fecundity without requiring species-specific terms (Fig. 3). That we did not find a significant interaction between density and species identity highlights the general ability of these five traits to describe multi-species fecundity without incorporating a species-specific response to the density gradient.

It is important to note that the changes in trait values observed here are primarily the result of phenotypic plasticity, in addition to genotype sorting due to differential mortality. Plasticity is widespread in nature and can impact demographic rates, life history, and species interactions [45]. Though trait-environment relationships are often conceptualized in terms of adaptation or species sorting, plasticity can also produce strong trait-environment relationships (see [46] for an example where light conditions determine leaf structure through plasticity). Zooplankton species are known to exhibit an exceptional range of adaptive plasticity [47]. Thus, predicting the response of zooplankton species to environmental change almost certainly requires that plasticity (and differences in plasticity among species or traits) be incorporated as a mechanism by which trait-environment relationships can develop [47]. Differences in plasticity across species and traits are likely common in nature and could explain some of the observed variation in trait-environment relationships, however, further study is necessary. Work with controlled experimental systems may be ideal for manipulating and measuring plasticity and genetic variation in order to place plastic changes in the context of trait-environmental relationships.

We tested two fundamental questions related to the utility of using relationships between traits and the environment to understand performance. We identified, consistent with other works (e.g., [10]), important explanatory or predictive relationships between traits and performance for multiple species. We found less support for the assumption that trait-environment relationships are general across taxonomic scales or among species. Hopefully these results will inform and improve the use of traits as a tool for predicting how changing environments and human impacts will affect species abundances and distributions in the future.

## Materials and Methods

### Experimental conditions

We used four daphniid species that are typically found in freshwater ponds and lakes: *Daphnia magna, Moina micrura, Ceriodaphnia dubia*, and *Simocephalus vetulus*. We established microcosms in 125 mL glass jars cultured under standard laboratory conditions with five density treatments per species, and each treatment was replicated at least 10 times. Microcosms were fed to maintain species-specific concentrations (cells/mL media) of *Chlamydomonas moewusii* three times weekly. From populations with standardized conditions, we collected pre-reproductive females, added them to 125 mL jars (1 female per jar), and allowed them to grow and reproduce. We selected 1-day old juveniles from their first brood to start the experimental microcosms, adding either 1, 2, 4, 6, or 8 same-age juveniles depending on the density treatment. Once established, we checked each microcosm daily (except on Sundays) and recorded demographic information including the number of juveniles born and adult mortality for each day. Juveniles were then removed, maintaining the initial treatment density; adults that died were not replaced since mortality is a meaningful outcome of high-density conditions. The experiments lasted for a single generation, and individuals were collected when they reached ~¾ of their average life expectancy. This length of time varied by species (*M. micrura* = 12 days, *C. dubia* = 16 days, *D. magna* = 20 days, *S. vetulus* = 30 days). Collecting individuals before they naturally senesce is essential, as deaths are unpredictable and decomposition occurs rapidly, making it difficult to measure traits accurately. Collected adults were euthanized with 90% ethanol, photographed, and then dried for 24h in a 60º C oven and weighed.

### Daphniid traits

In order to understand the independent contribution of each morphological trait (body length, 2nd antenna, and eye diameter) to a functional relationship between the density gradient and performance, beyond their strong correlations with total body mass, we detrended these traits for body mass (dry mass, mg, see Table S1 for values by species). We fit an allometric model to each morphological trait (per species) as a function of body mass:

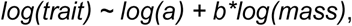

where *a* is a normalization constant and *b* the scaling term (e.g.[48]). For each individual measurement, the difference between the observed and predicted trait value was used as the new body mass-detrended trait value. To calculate individual relative growth rate (RGR), we used independent data which included, for all species, biomass measurements for ~20 replicate individuals per day, for a 20-day time series. We fit a Weibull function to each species’ data, which describes body mass as a saturating function of time, and retained the species-specific estimates of minimum individual mass. Then, for each experimental replicate, we parameterized a Weibull function with the species-specific estimates of minimum body mass, set maximum intraspecific body mass as the maximum size observed in that respective density treatment, and then used observed body mass to solve for the remaining variable – RGR of that individual. For microcosms with multiple surviving individuals, we used the average trait values for the replicate. Finally, we used fecundity as the measure of performance, calculated as the average per capita rate of juvenile production (juveniles/adult/day). For each treatment with more than one adult individual, we divided the total number of juveniles born by the weighted mean number of adults present in a jar over the course of the experiment.

### Statistical Analyses

We wanted to know which traits interact with the conspecific density gradient to determine individual performance. To improve normality, we log-transformed length, 2nd antenna, eye diameter, and RGR, and then standardized all traits by unit variance. Standardization was applied within-species (intraspecific) for single species analyses, and across-species for the multi-species (interspecific) analyses. We used the R package *piecewiseSEM* (version 2.0.2, [49]) to fit structural equation models (SEMs). We first created the full SEM which incorporated all possible links between density and the measured traits (body length, 2nd antenna, eye diameter, age at first reproduction, and relative growth rate), and between these traits and fecundity (Fig. 2a). Each path in the SEM was represented by a single model. Conspecific density was treated as an exogenous variable, and potential paths between traits and density were fit with linear models. For the intraspecific model, individual fecundity was scaled by the maximum fecundity for each species and then log-transformed to increase normality. In the interspecific SEM, fecundity was scaled to the maximum fecundity value across species, then log transformed. We initially considered both linear and quadratic forms and used AIC to determine if a quadratic model should be retained; as we found no support for retaining the quadratic terms, all SEM functions are linear.

We evaluated all unspecified claims for non-independence using tests of directed separation and ensured that all significant pathways were included. Goodness-of-fit was assessed using these tests of directed separation, which gives a Fisher’s *C* statistic that is *X*2 distributed. Large *p*-values (> 0.05) associated with Fisher’s *C* indicate that the model represents the data well. We also fit the data to a second model in which we explicitly tested for a direct link between fecundity and density, however, in every case, this model was not favored over the model in which these parameters are treated as correlated errors (deltaAIC ≤ 2). After fitting the full model per species, we also fit a nested version containing only the significant links identified in the full model to confirm, using AIC, that the full model is not more likely than the reduced model, given the data. We defined significant trait-environment relationships as those for which both the path from density to the trait, and from the trait to fecundity were significant (*p* < 0.05). For all species, the nested SEM was supported over the full model (deltaAIC >> 2); however, we report the results of the full models so that direct comparisons of path coefficients and model structure can be made across species. Nested model results are in the Supplementary Materials (Table S2).

Additionally, to ask whether there are general changes in trait values associated with the density gradient across all species, we applied a permutational MANOVA (using *RRPP*, [50]) of the form:

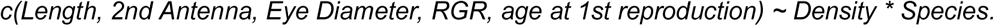

All analyses were performed in R (version 3.5.2, R Core Team 2018).

## Acknowledgments

Many thanks to B. Enquist, D. Goldberg, G. Legault, A. Rolhauser, and to the members of the 3rd Floor Ecology group at UNC for their feedback on various aspects and versions of this project. Thank you as well to all lab assistants, including E. Kremer, E. Feldmann, S. Charlino, Z. Yu, N. Nair, S. Long, K. May, and L. Stiller, without whom this work would be impossible.

