## Supplementary Materials for "Trait-environment relationships are predictive, but not general across species"

**Supplemental Information for Frances, Barber and Tucker (2020)**

**Supplemental Methods.** Source cultures of each zooplankton species were originally obtained from multiple sources, including biological suppliers (Carolina Biological, Burlington, N.C.), aquaculture suppliers (Sachs Systems Aquaculture, St. Augustine, F.L.), and ponds local to the University of North Carolina. Source cultures have been maintained in the lab for more than 1.5 years, and have been observed to undergo multiple asexual and sexual cycles during this time. For experimental purposes, subsampled populations are established from the source culture, and these are maintained for multiple generations (at least 4) before use to ensure the absence of sexual individuals; further sexual reproduction is avoided by limiting population densities (Gerber *et al* 2018) These pre-experiment populations were cultured at conditions (food, media, temperature) identical to those to be used in the experiment. Microcosms for all species use 100mL of modified COMBO media (originally from Kluttgen *et al* 1994) in 125 mL wide-mouthed glass jars. These microcosms are cultured under standardized laboratory conditions (23º C ± 0.5; photoperiod 14:10 L:D). Microcosms are fed to maintain species-specific concentrations (cells/mL media) of *Chlamydomonas moewusii*; feedings occur on a Monday/Wednesday/ Friday schedule.

**Feeding concentrations.** Initial *Chlamydomonas moewusii* samples (CC-2529 C. moewusii LM74) were obtained from the Chlamydomonas Resource Center (CRC). *Chlamydomonas moewusii* is cultured semi-continuously in the lab in 2L flasks containing algal media (protocol available on request), to which a vitamin supplement containing additional micronutrients is added after 2 days of culture. Flasks are collected after 1-1.5 weeks of growth, and to concentrate the algal cells, the media is centrifuged at 3500 rpm (acceleration 8) for 10 minutes and the supernatant is poured off. The concentration is measured using a spectrophotometer at 680 nm, using a species-specific calibration curve to calculate the concentration.
Feeding concentrations for each species were determined as:

*Daphnia magna:* $4.00 \times 10_7$ cells/L
*Simocephalus vetulus:* $7.50 \times 10_6$ cells/L
*Ceriodaphnia dubia:* $7.00 \times 10_6$ cells/L
*Moina micrura:* $3.00 \times 10_7$ cells/L

**Experimental methods.** We established microcosms for five density treatments per species, with each treatment replicated at least 10 times. Juvenile females were selected from the pre-experiment populations and added individually to 125 mL experimental jars and allowed to grow and reproduce. From this maternal generation we used 1-day old juveniles from the mother's first brood to start the experimental replicates. This approach helps to minimize variation from maternal effects, brood effects, and age variation in the microcosms. When starting a microcosm, 2x the usual amount of food was added. Once established, each microcosm was checked daily (except on Sundays; juveniles born on Sundays were aged by size and recorded appropriately the next day), and daily demographic information, including the number of juveniles born and adult mortality, was recorded. Juveniles were then removed, maintaining the initial treatment density; adults that died were not replaced since mortality is a meaningful outcome of high-density conditions. The experiments lasted for a single generation, and individuals were collected when they reached ~¾ of their average life expectancy. This length of time varied by species (*M. micrura* = 12 days, *C. dubia* = 16 days, *D. magna* = 20 days, *S. vetulus* = 30 days). Collecting individuals before they die is essential, as deaths are unpredictable and decomposition occurs rapidly, making it difficult to measure traits accurately. Adults that survived were euthanized with ethanol and then photographed with a CX43 microscope with LC30 digital camera and cellSens

Entry imaging software (version 2.1, Olympus), using objectives 2X-10X. Individuals were then dried at 60º C for 24 hours and massed on a microbalance (XPR2, Mettler Toledo, accurate to 1 ug)].

### Supplemental Figures

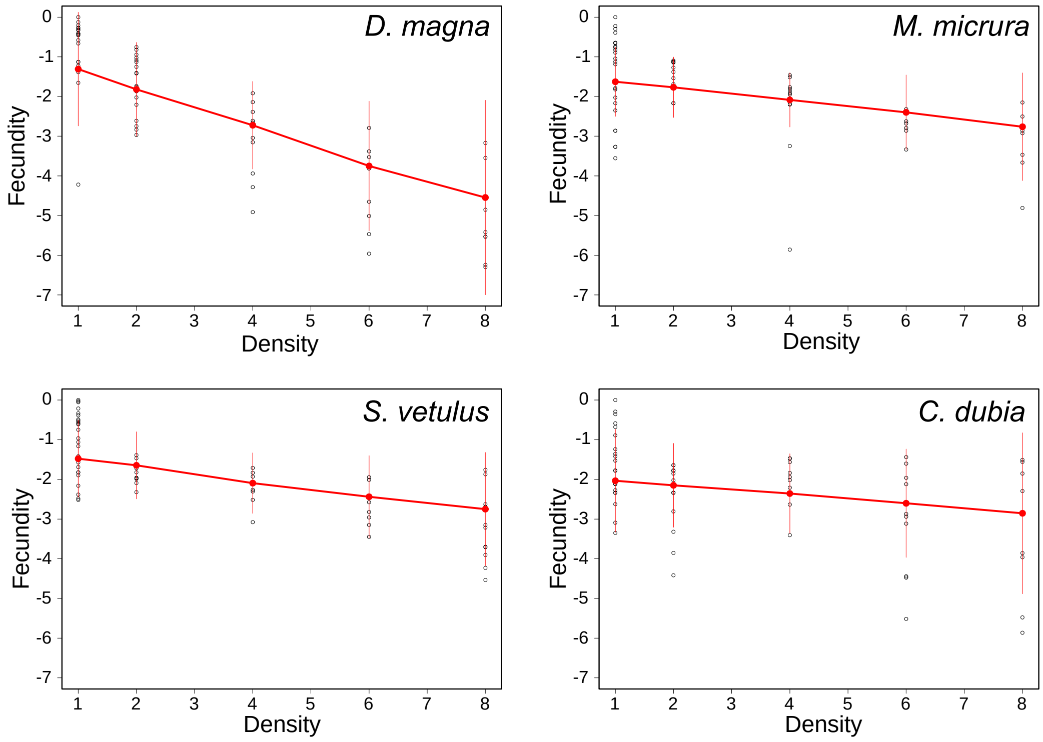


**Figure S1**. Model fit in red, showing predicted fecundity per density treatment for each species (*D. magna, M. micrura, S. vetulus,* and *C. dubia*). We calculated the mean and standard error of each trait for a given density treatment level and used these to define a normal distribution for each trait combination. We drew randomly from these distributions and used the trait values to calculate fecundity. This procedure was repeated 1000 times for each density. Fecundity is a rate of juveniles produced per adult per day, scaled by the max fecundity within each species, and then log-transformed. Error bars are 95% confidence intervals.

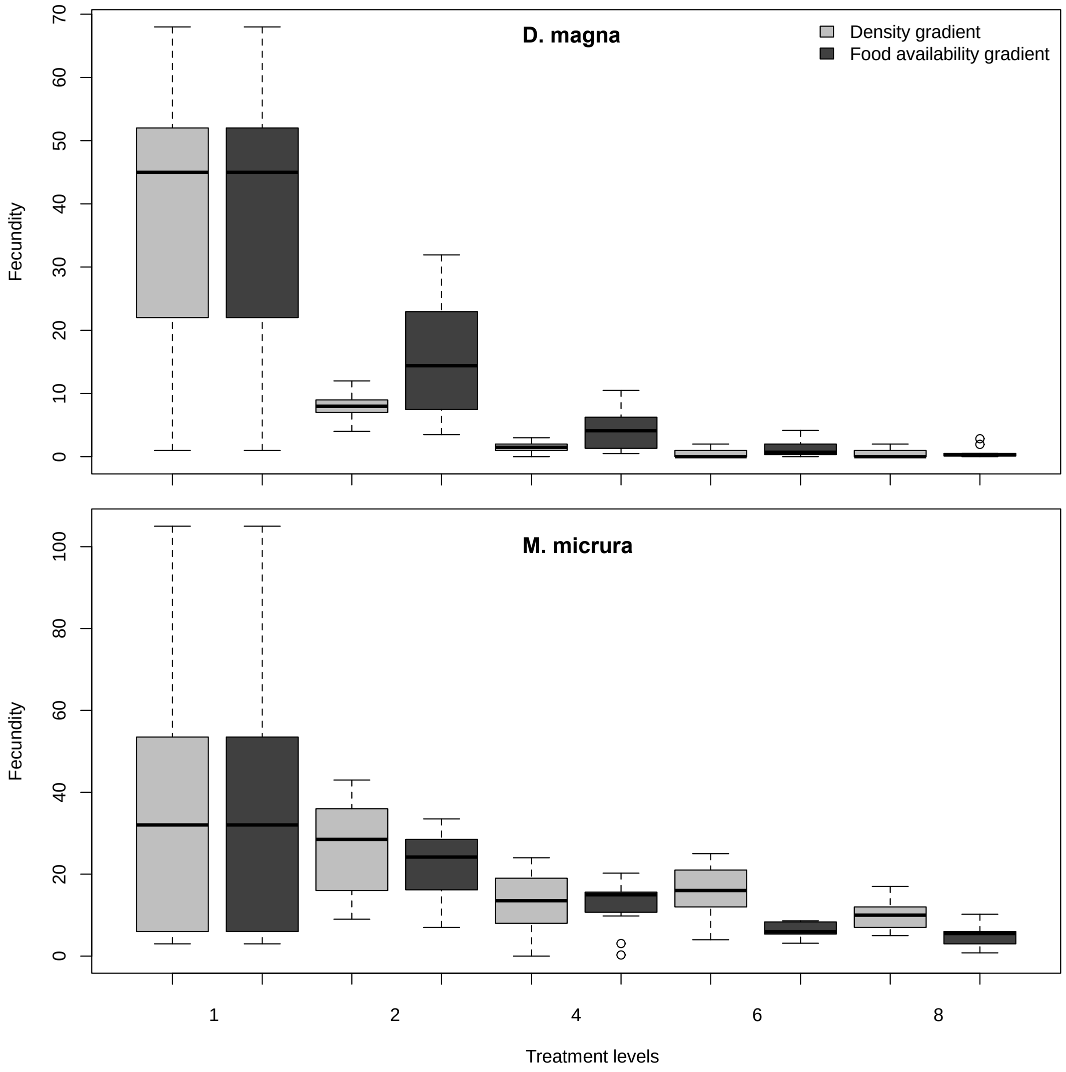


**Figure S2**. Comparing fecundity at each density level from the present experiment (light gray shading) to the food availability experiment (dark shading). In the additional study, algae concentrations were matched to the gradient of food available in the original experiment, but we fixed the number of individuals to one. There was no significant difference in fecundity from the present experiment to the food limitation study, indicating that food limitation is the proximate mechanism driving changes in performance and that large (*D. magna*) and small (*M. micrura*) species experience the density treatment in the same way ($p$ = 0.826 and $p$ = 0.490, respectively).

**Table S1.** Traits used for analyses. Includes a list of traits measured for each individual daphniid, a description of the measurement and the associated units, justification or relevance of the trait for daphniid ecology, and summary values per species.

| Trait | Description | Justification | Value (mean) | Units |
|---|---|---|---|---|
| **Body mass** | Dry weight per individual. | "Master trait". Associated with other morphological and physiological measures (Kiorboe et al 2014). | *D. magna* = 0.17<br>*C. dubia* = 0.01<br>*M. micrura* = 0.02<br>*S. vetulus* = 0.04 | mg |
| **Length** | Total body length, distance from top of head (above eye), to start of apical spine (if present). See Figure 1a. | Potential relationship to multiple ecological measures including predation risk, brood pouch size, vital rates, and patterns of energy allocation (Litchman et al. 2013; Boersma 1998). | *D. magna* = 3.00<br>*C. dubia* = 0.81<br>*M. micrura* = 1.30<br>*S. vetulus* = 1.50 | mm |
| **Eye diameter** | Diameter of compound eye. See Figure 1b. | Compound eyes require expensive materials during development, but are associated with increased fitness; reflects changes in resource allocation under stress (Boersma 1998; Brandon et al. 2015). | *D. magna* = 0.18<br>*C. dubia* = 0.07<br>*M. micrura* = 0.11<br>*S. vetulus* = 0.08 | mm |
| **2nd antenna** | Length of the main segment of the 2nd antenna. See Figure 1c. | Associated with individual movement and filter feeding efficiency (Dodson et al 1997; Ebert 2005). | *D. magna* = 0.66<br>*C. dubia* = 0.18<br>*M. micrura* = 0.34<br>*S. vetulus* = 0.24 | mm |
| **Age at first reproduction** | Individual age at the production of their first brood. | Life history character with known tradeoffs with other life history measures such as lifespan and total reproductive output (Harvey and Zammuto 1985; Vanni & Lampert 1992). | *D. magna* = 11.4<br>*C. dubia* = 9.3<br>*M. micrura* = 4.7<br>*S. vetulus* = 11.8 | day |
| **Relative growth rate** | Proportional rate of increase in mass through time. | Reflects energy allocation to growth. May be associated with higher reproductive ability later in time (Litchman et al. 2013). | *D. magna* = 0.04<br>*C. dubia* = 0.03<br>*M. micrura* = 0.04<br>*S. vetulus* = 0.01 | mg/day |

**Tables S2**

***D. magna***

### Full model AIC: 60.775

Global goodness-of-fit:

Fisher's C = 16.775 with P-value = 0.268 and on 14 degrees of freedom

Coefficients:

| Response | Predictor | DF | Crit.Value | P.Value | Std.Estimate |
|---|---|---|---|---|---|
| Length | Density | 62 | -1.4507 | 0.1519 | -0.1812 |
| Eye diameter | Density | 62 | -0.2716 | 0.7868 | -0.0345 |
| 2nd antenna | Density | 62 | -1.5618 | 0.1234 | -0.1946 |
| RGR | Density | 62 | -8.2249 | 0 | -0.7223 |
| First brood | Density | 62 | 2.8106 | 0.0066 | 0.3362 |
| Fecundity | RGR | 58 | 9.3487 | 0 | 0.6456 |
| Fecundity | Length | 58 | 1.5139 | 0.1355 | 0.1613 |
| Fecundity | 2nd antenna | 58 | 2.3956 | 0.0198 | 0.2012 |
| Fecundity | Eye diameter | 58 | -0.7633 | 0.4484 | -0.0655 |
| Fecundity | First brood | 58 | -4.5907 | 0 | -0.3306 |
| ~~Fecundity | ~~Density | 64 | -3.6242 | 0.0003 | -0.4209 |
| ~~Eye diameter | ~~Length | 64 | 6.1216 | 0 | 0.6169 |
| ~~2nd antenna | ~~Length | 64 | 5.3203 | 0 | 0.563 |
| ~~RGR | ~~2nd antenna | 64 | -2.0384 | 0.0229 | -0.2525 |

| Response | method | R.squared |
|---|---|---|
| Length | none | 0.03 |
| Eye diameter | none | 0 |
| 2nd antenna | none | 0.04 |
| RGR | none | 0.52 |
| First brood | none | 0.11 |
| Fecundity | nagelkerke | 0.95 |

### Nested model AIC: 37.793

Fisher's C = 13.793 with P-value = 0.465 and on 14 degrees of freedom

| Response | Predictor | DF | Crit.Value | P.Value | Std.Estimate |
|---|---|---|---|---|---|
| RGR | Density | 62 | -8.2249 | 0 | -0.7223 |
| First brood | Density | 62 | 2.8106 | 0.0066 | 0.3362 |
| Fecundity | Length | 59 | 1.3282 | 0.1892 | 0.1115 |
| Fecundity | 2nd antenna | 59 | 2.6523 | 0.0103 | 0.216 |
| Fecundity | RGR | 59 | 9.3529 | 0 | 0.6425 |
| Fecundity | First brood | 59 | -4.686 | 0 | -0.3351 |
| ~~Fecundity | ~~Density | 64 | -3.699 | 0.0002 | -0.428 |
| ~~RGR | ~~2nd antenna | 64 | -1.9968 | 0.0252 | -0.2477 |

| Response | method | R.squared |
|---|---|---|
| RGR | none | 0.52 |
| First brood | none | 0.11 |
| Fecundity | nagelkerke | 0.95 |

***M. micrura***

**Full model AIC: 53.757**

Global goodness-of-fit:

Fisher's C = 9.757 with P-value = 0.637 and on 12 degrees of freedom

| Response | Predictor | DF | Crit.Value | P.Value | Std.Estimate |
|---|---|---|---|---|---|
| Length | Density | 61 | -4.178 | 0.0001 | -0.4717 |
| Eye diameter | Density | 61 | -2.9168 | 0.0049 | -0.3499 |
| 2nd antenna | Density | 61 | -1.5433 | 0.1279 | -0.1939 |
| RGR | Density | 61 | -2.4895 | 0.0155 | -0.3037 |
| First brood | Density | 61 | -1.1188 | 0.2676 | -0.1418 |
| Fecundity | RGR | 57 | 2.9179 | 0.005 | 0.2819 |
| Fecundity | Length | 57 | 3.2693 | 0.0018 | 0.4 |
| Fecundity | 2nd antenna | 57 | -1.3362 | 0.1868 | -0.1479 |
| Fecundity | Eye diameter | 57 | 3.4128 | 0.0012 | 0.3768 |
| Fecundity | First brood | 57 | -1.228 | 0.2245 | -0.1225 |
| ~~Fecundity | ~~Density | 63 | -2.358 | 0.0108 | -0.2912 |
| ~~Eye diameter | ~~Length | 63 | 2.5521 | 0.0066 | 0.3129 |
| ~~2nd antenna | ~~Length | 63 | 3.8471 | 0.0001 | 0.4448 |
| ~~RGR | ~~Length | 63 | -2.3307 | 0.0116 | -0.2881 |
| ~~First brood | ~~Eye diameter | 63 | -2.4045 | 0.0096 | -0.2965 |

| Response | method | R.squared |
|---|---|---|
| Length | none | 0.22 |
| Eye diameter | none | 0.12 |
| 2nd antenna | none | 0.04 |
| RGR | none | 0.09 |
| First brood | none | 0.02 |
| Fecundity | nagelkerke | 0.62 |

**Nested model AIC: 41.324**

Fisher's C = 13.324 with P-value = 0.501 and on 14 degrees of freedom

| Response | Predictor | DF | Crit.Value | P.Value | Std.Estimate |
|---|---|---|---|---|---|
| Length | Density | 61 | -4.178 | 0.0001 | -0.4717 |
| Eye diameter | Density | 61 | -2.9168 | 0.0049 | -0.3499 |
| RGR | Density | 61 | -2.4895 | 0.0155 | -0.3037 |
| Fecundity | RGR | 59 | 2.9315 | 0.0048 | 0.285 |
| Fecundity | Length | 59 | 2.8872 | 0.0054 | 0.3098 |
| Fecundity | Eye diameter | 59 | 3.995 | 0.0002 | 0.4267 |
| ~~Fecundity | ~~Density | 63 | -2.038 | 0.023 | -0.2544 |
| ~~Eye diameter | ~~First brood | 63 | -2.3779 | 0.0103 | -0.2935 |
| ~~Length | ~~2nd antenna | 63 | 3.7568 | 0.0002 | 0.4364 |
| ~~Eye diameter | ~~Length | 63 | 2.5521 | 0.0066 | 0.3129 |
| ~~RGR | ~~Length | 63 | -2.3307 | 0.0116 | -0.2881 |

| Response | method | R.squared |
|---|---|---|
| Length | none | 0.22 |
| Eye diameter | none | 0.12 |
| RGR | none | 0.09 |
| Fecundity | nagelkerke | 0.59 |

### *S. vetulus*

#### Full model AIC: 58.196

Global
goodness-of-fit:
Fisher's C = 14.196 with P-value = 0.288
and on 12 degrees of freedom

| Response | Predictor | DF | Crit.Value | P.Value | Std.Estimate |
|---|---|---|---|---|---|
| Length | Density | 62 | -3.8794 | 0 | -0.442 |
| Eye diameter | Density | 62 | -3.7491 | 0 | -0.4299 |
| 2nd antenna | Density | 62 | -0.3237 | 0.75 | -0.0411 |
| RGR | Density | 62 | -2.7727 | 0.01 | -0.3321 |
| First brood | Density | 62 | 1.73 | 0.09 | 0.2146 |
| Fecundity | RGR | 58 | 7.0287 | 0 | 0.485 |
| Fecundity | Length | 58 | 5.9407 | 0 | 0.4877 |
| Fecundity | 2nd antenna | 58 | -2.229 | 0.03 | -0.1677 |
| Fecundity | Eye diameter | 58 | 2.2424 | 0.03 | 0.1712 |
| Fecundity | First brood | 58 | -3.0585 | 0 | -0.2305 |
| ~~Fecundity | ~~Density | 64 | -4.658 | 0 | -0.5122 |
| ~~Eye diameter | ~~Length | 64 | 2.4049 | 0.01 | 0.2943 |
| ~~2nd antenna | ~~Length | 64 | 3.6698 | 0 | 0.4253 |
| ~~First brood | ~~Length | 64 | -2.9408 | 0 | -0.3524 |
| ~~2nd antenna | ~~Eye diameter | 64 | 3.4434 | 0 | 0.4034 |

| Response | method | R.squared |
|---|---|---|
| Length | none | 0.2 |
| Eye diameter | none | 0.18 |
| 2nd antenna | none | 0 |
| RGR | none | 0.11 |
| First brood | none | 0.05 |
| Fecundity | nagelkerke | 0.83 |

#### Nested model AIC: 42.779

Fisher's C = 10.779 with P-value = 0.375 and on 10
degrees of freedom

| Response | Predictor | DF | Crit.Value | P.Value | Std.Estimate |
|---|---|---|---|---|---|
| Length | Density | 62 | -3.8794 | 0.0003 | -0.442 |
| Eye diameter | Density | 62 | -3.7491 | 0.0004 | -0.4299 |
| RGR | Density | 62 | -2.7727 | 0.0073 | -0.3321 |
| Fecundity | Eye diameter | 58 | 2.2424 | 0.0288 | 0.1712 |
| Fecundity | 2nd antenna | 58 | -2.229 | 0.0297 | -0.1677 |
| Fecundity | RGR | 58 | 7.0287 | 0 | 0.485 |
| Fecundity | Length | 58 | 5.9407 | 0 | 0.4877 |
| Fecundity | First brood | 58 | -3.0585 | 0.0034 | -0.2305 |
| ~~Fecundity | ~~Density | 64 | -4.658 | 0 | -0.5122 |
| ~~Length | ~~First brood | 64 | -2.863 | 0.0029 | -0.3442 |
| ~~Length | ~~2nd antenna | 64 | 3.666 | 0.0003 | 0.4249 |
| ~~Eye diameter | ~~2nd antenna | 64 | 3.4399 | 0.0005 | 0.4031 |
| ~~Eye diameter | ~~Length | 64 | 2.4049 | 0.0096 | 0.2943 |

| Response | method | R.squared |
|---|---|---|
| Length | none | 0.2 |
| Eye diameter | none | 0.18 |
| RGR | none | 0.11 |
| Fecundity | nagelkerke | 0.83 |

***C. dubia***

### Full model AIC: 48.285

Global goodness-of-fit:
Fisher's C = 4.285 with P-value = 0.993 and on 14 degrees of freedom

| Response | Predictor | DF | Crit.Value | P.Value | Std.Estimate |
|---|---|---|---|---|---|
| Length | Density | 61 | -1.4589 | 0.1497 | -0.1836 |
| Eye diameter | Density | 61 | -0.9288 | 0.3566 | -0.1181 |
| 2nd antenna | Density | 61 | 1.0171 | 0.3131 | 0.1291 |
| RGR | Density | 61 | -0.5427 | 0.5893 | -0.0693 |
| First brood | Density | 61 | 1.9253 | 0.0589 | 0.2393 |
| Fecundity | RGR | 57 | 1.3768 | 0.1739 | 0.1143 |
| Fecundity | Length | 57 | 2.697 | 0.0092 | 0.318 |
| Fecundity | 2nd antenna | 57 | -2.391 | 0.0201 | -0.2503 |
| Fecundity | Eye diameter | 57 | -0.9262 | 0.3582 | -0.1076 |
| Fecundity | First brood | 57 | -7.749 | 0 | -0.6631 |
| ~~Fecundity | ~~Density | 63 | -2.7579 | 0.0038 | -0.3354 |
| ~~Eye diameter | ~~Length | 63 | 6.6281 | 0 | 0.6502 |
| ~~2nd antenna | ~~Length | 63 | 5.5112 | 0 | 0.5797 |
| ~~2nd antenna | ~~Eye diameter | 63 | 5.505 | 0 | 0.5793 |

| Response | method | R.squared |
|---|---|---|
| Length | none | 0.03 |
| Eye diameter | none | 0.01 |
| 2nd antenna | none | 0.02 |
| RGR | none | 0 |
| First brood | none | 0.06 |
| Fecundity | nagelkerke | 0.77 |

### Nested model AIC: 15.263

Fisher's C = 5.263 with P-value = 0.261 and on 4 degrees of freedom

| Response | Predictor | DF | Crit.Value | P.Value | Std.Estimate |
|---|---|---|---|---|---|
| Fecundity | Length | 59 | 2.5549 | 0.0132 | 0.2608 |
| Fecundity | 2nd antenna | 59 | -2.755 | 0.0078 | -0.2762 |
| Fecundity | First brood | 59 | -7.8178 | 0 | -0.6711 |
| ~~Fecundity | ~~Density | 63 | -2.695 | 0.0046 | -0.3286 |

| Response | method | R.squared |
|---|---|---|
| Fecundity | nagelkerke | 0.75 |

**All Species (Interspecific model)**

### Full model AIC: 50.505

Global goodness-of-fit:

Fisher's C = 6.505 with P-value = 0.591 and on 8 degrees of freedom

| Response | Predictor | DF | Crit.Value | P.Value | Std.Estimate |
|---|---|---|---|---|---|
| Length | Density | 252 | -5.4801 | 0 | -0.3263 |
| Eye diameter | Density | 252 | -3.7176 | 0.0002 | -0.228 |
| RGR | Density | 252 | -5.3582 | 0 | -0.3198 |
| 2nd antenna | Density | 252 | -1.2904 | 0.1981 | -0.081 |
| First brood | Density | 252 | 2.605 | 0.0097 | 0.1619 |
| Fecundity | Length | 248 | 4.783 | 0 | 0.2879 |
| Fecundity | 2nd antenna | 248 | -1.4783 | 0.1406 | -0.078 |
| Fecundity | Eye diameter | 248 | 0.6281 | 0.5305 | 0.034 |
| Fecundity | First brood | 248 | -9.4426 | 0 | -0.4623 |
| Fecundity | RGR | 248 | 5.5293 | 0 | 0.2666 |
| ~~Fecundity | ~~Density | 254 | 0.8086 | 0.2098 | 0.051 |
| ~~Eye diameter | ~~Length | 254 | 9.015 | 0 | 0.4946 |
| ~~2nd antenna | ~~Length | 254 | 9.0028 | 0 | 0.4941 |
| ~~First brood | ~~Length | 254 | -2.004 | 0.0231 | -0.1255 |
| ~~2nd antenna | ~~Eye diameter | 254 | 4.7379 | 0 | 0.2865 |
| ~~RGR | ~~Length | 254 | -2.151 | 0.0162 | -0.1345 |
| ~~First brood | ~~RGR | 254 | -4.3836 | 0 | -0.2667 |

| Response | method | R.squared |
|---|---|---|
| Length | none | 0.11 |
| Eye diameter | none | 0.05 |
| RGR | none | 0.1 |
| 2nd antenna | none | 0.01 |
| First brood | none | 0.03 |
| Fecundity | nagelkerke | 0.72 |

### Nested model AIC: 44.335

Fisher's C = 10.335 with P-value = 0.587 and on 12 degrees of freedom

| Response | Predictor | DF | Crit.Value | P.Value | Std.Estimate |
|---|---|---|---|---|---|
| Length | Density | 252 | -5.4801 | 0 | -0.3263 |
| Eye diameter | Density | 252 | -3.7176 | 0.0002 | -0.228 |
| RGR | Density | 252 | -5.3582 | 0 | -0.3198 |
| First brood | Density | 252 | 2.605 | 0.0097 | 0.1619 |
| Fecundity | Length | 250 | 5.7175 | 0 | 0.2669 |
| Fecundity | First brood | 250 | -9.5091 | 0 | -0.4655 |
| Fecundity | RGR | 250 | 5.5183 | 0 | 0.2663 |
| ~~Fecundity | ~~Density | 254 | 0.8455 | 0.1993 | 0.0533 |
| ~~Length | ~~2nd antenna | 254 | 0.7907 | 0.2149 | 0.0498 |
| ~~Eye diameter | ~~2nd antenna | 254 | 1.1613 | 0.1233 | 0.0731 |
| ~~Eye diameter | ~~Length | 254 | 9.015 | 0 | 0.4946 |
| ~~RGR | ~~Length | 254 | -2.151 | 0.0162 | -0.1345 |
| ~~First brood | ~~Length | 254 | -2.004 | 0.0231 | -0.1255 |
| ~~First brood | ~~RGR | 254 | -4.3836 | 0 | -0.2667 |

| Response | method | R.squared |
|---|---|---|
| Length | none | 0.11 |
| Eye diameter | none | 0.05 |
| RGR | none | 0.1 |
| First brood | none | 0.03 |
| Fecundity | nagelkerke | 0.71 |

**Table S3.** Multivariate analysis of variance.
Formula:
*c(Length, 2nd Antenna, Eye Diameter, RGR, Age at first reproduction) ~ Density * Species*

| **Term** | **Df** | **SS** | **MS** | $R^2$ | $F$ | **Z** | $p$ |
|---|---|---|---|---|---|---|---|
| **Conspecific density** | 1 | 149.8 | 149.80 | 0.02835 | 12.2813 | 2.1127 | **0.00050** |
| **Species** | 3 | 2040.7 | 680.24 | 0.38618 | 55.7709 | 4.8279 | **9.999e-05** |
| **Density * Species** | 3 | 93.4 | 31.13 | 0.01767 | 2.5525 | 1.4221 | 0.05119 |
| **Residuals** | 246 | 3000.5 | 12.20 | 0.56780 | | | |
| **Total** | 253 | 5284.4 | | | | | |

**Tables S4. Food manipulation experiment.**
Comparison of experimental gradients in which conspecific density is manipulated vs. those in which only food amount available is manipulated and their effects on fecundity.

a. *Daphnia magna*

| **Term** | **Estimate** | **St. Error** | **t value** | ***p*** | |
|---|---|---|---|---|---|
| **Intercept** | 34.8835 | 3.2338 | 10.787 | **< 2e-16** | |
| **Gradient level** | -5.4366 | 0.7315 | -7.432 | **1.57e-11** | |
| **Gradient type** | -0.2017 | 4.3913 | -0.046 | 0.963 | |
| **Gradient level * Gradient type** | 0.2240 | 1.0140 | 0.221 | 0.826 | $R_2$: 0.4608 |

b. *Moina micrura*

| **Term** | **Estimate** | **St. Error** | **t value** | ***p*** | |
|---|---|---|---|---|---|
| **Intercept** | 34.7594 | 3.7114 | 9.366 | **< 2e-16** | |
| **Gradient level** | -3.4715 | 0.8913 | -3.895 | **0.000154** | |
| **Gradient type** | 0.3300 | 1.2877 | 0.063 | 0.950072 | |
| **Gradient level * Gradient type** | -0.8906 | 1.0140 | -0.692 | 0.490315 | $R_2$: 0.1991 |
